# Neonatal regulatory T cells promote alveologenesis by restraining neutrophil-mediated inflammation

**DOI:** 10.64898/2026.03.24.714021

**Authors:** Monica Sen, Prudence Pokwai Lui, Hafsah Aziz, Jessie Z. Xu, Niwa Ali

## Abstract

Alveologenesis during early postnatal life requires tight coordination between epithelial differentiation and immune regulation, yet how immune cells contribute to this process remains unclear. Regulatory T cells (Tregs) are established mediators of immune homeostasis and tissue repair in adult lung injury, but their role in lung development is unknown. Here, we identify a transient wave of highly proliferative, activated Tregs that accumulates in the neonatal lung during an early postnatal window. Using inducible Treg ablation, we show that loss of this neonatal Treg population disrupts alveologenesis, resulting in enlarged airspaces, and persistent structural abnormalities later in life. Treg depletion also induces interferon-associated inflammatory programmes, promotes neutrophil accumulation, and is accompanied by a sustained imbalance in alveolar epithelial populations. Notably, neutrophil depletion partially rescues both epithelial composition and alveolar structure, identifying neutrophils as key downstream effectors of Treg-mediated regulation. Together, these findings show that neonatal Tregs are required for normal alveologenesis by restraining neutrophil-driven inflammation and preserving epithelial balance. Our study reveals a previously unappreciated role for immune regulation in lung organogenesis.

## Introduction

Regulatory T cells (Tregs) are best known for restraining excessive immune responses and maintaining peripheral tolerance. However, it is increasingly clear that Tregs also acquire tissue-adapted functions that extend beyond classical immune suppression and contribute directly to tissue homeostasis, repair and remodelling^1–3^. Although a substantial proportion of Tregs reside in lymphoid organs, distinct Treg populations are established within non-lymphoid tissues^4^, where they adopt specialised phenotypes shaped by local environmental cues and perform context-specific functions^5–8^.

The lung undergoes profound structural remodelling during the early postnatal period, when alveolar formation requires tight coordination between epithelial differentiation, septation and inflammatory control^9,10^. Disruption of this balance during neonatal life can have lasting consequences for lung architecture and function. In adult LPS-induced acute lung injury models, the loss of Tregs delays the resolution of lung inflammation^11,12^, accompanied by upregulation of neutrophils and alveolar macrophages abundance^13^, but also delayed neutrophil clearance^11^ and significantly inhibits EpCam+ lung epithelial cells proliferation^12^, positioning lung Tregs as key coordinators linking immune resolution with tissue remodelling in adult lung regeneration. Yet, whether Tregs contribute to normal lung development during the neonatal period remains unknown.

We previously showed that an early wave of skin-seeding Tregs is required to establish later-life pigmentation^8^, suggesting that tissue Tregs can play instructive roles during postnatal organ development. We therefore hypothesised that a similarly timed population of neonatal lung Tregs might support alveolar development by restraining inappropriate inflammation during this critical developmental window. Here, we identify a transient wave of highly proliferative and activated Tregs that accumulates in the early postnatal lung. Using neonatal Treg depletion, transcriptional profiling and neutrophil rescue experiments, we show that these cells are required to preserve epithelial balance and proper alveolarisation by limiting early-life inflammatory programmes and neutrophil accumulation. Together, these findings reveal a previously unappreciated role for Tregs in organogenesis and identify neonatal lung Tregs as key regulators of postnatal lung development.

## Results

### A transient wave of activated Tregs characterises the early postnatal lung

To define the temporal dynamics of lung regulatory T cells (Tregs), we profiled CD4^+^Foxp3^+^ cells from birth (gating strategy in SFig1A). Tregs were detectable as early as postnatal day 3 (P3), peaked at P5, and declined progressively thereafter (Fig1A&B), consistent with previous reports of higher Treg abundance in neonatal compared to adult lungs^14^. This early accumulation resulted in a significantly higher proportion of Tregs among CD4^+^ T cells during the first postnatal week compared to adulthood.

**Figure 1.**
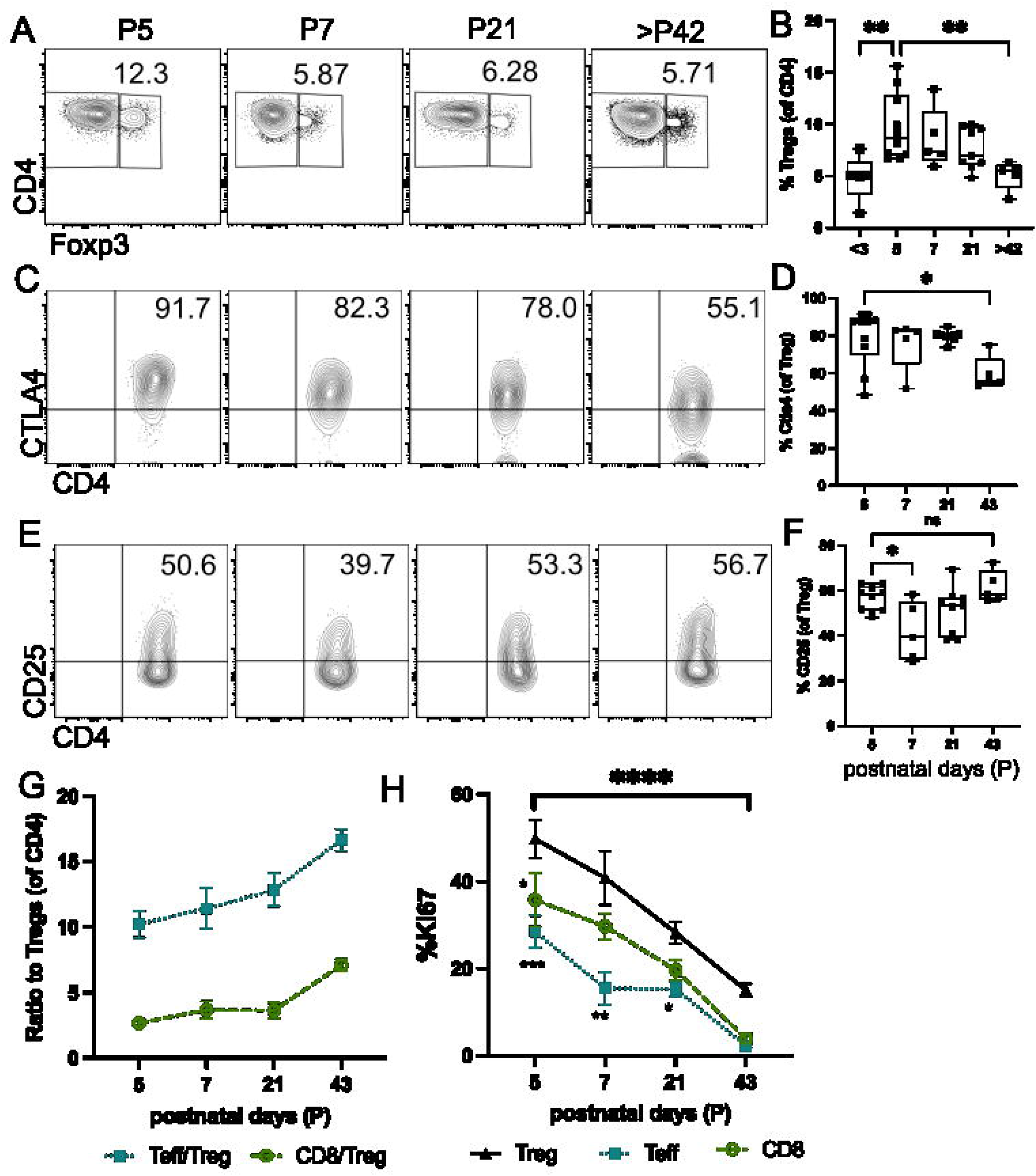
A transient wave of activated, proliferating Tregs marks the early postnatal lung. (A) Representative flow cytometry plots showing Foxp3+ Tregs abundance at postnatal day (P)5, P7, P21, and >P42. (B) Quantification of Tregs across postnatal time points. Representative flow cytometry plots and quantification of (C&D) CTLA-4 and (E&F) CD25 expression within Tregs at the indicated time points. (G) Ratio of effector T cells (Teffs, in blue square) and CD8+ T cells (in green circle) to Tregs across postnatal days. (H) %Ki67 of Treg, Teffs, and CD8^+^ T cells over indicated ages. Data are presented as mean ± SEM. Each point represents an individual sample, pooled from 4 experiments with n =5-10 for each timepoint. Statistical significance was determined using one-way ANOVA. *p < 0.05, **p < 0.01, \*\*\**p < 0*.*0001*.

At P5, a higher proportion of lung Tregs expressed CTLA-4 (Fig1C&D) and CD25 (Fig1E&F), indicative of an activated phenotype. In contrast, the ratio of conventional CD4^+^ T cells (Teff) and CD8^+^ T cells to Tregs increased with age (Fig1G), reflecting a progressive shift away from a Treg-enriched environment.

Lung Tregs at P5 also exhibited markedly elevated Ki67 expression compared to Teff and CD8^+^ T cells (Fig1H), indicating a strong proliferative burst. This proliferative activity declined sharply thereafter, from 49.8% at P5 to 15% in adulthood. This suggests Treg expansion is temporally restricted to this early postnatal window. Together, these data identify a transient wave of highly proliferative, activated Tregs that characterises the early postnatal lung.

### Neonatal Tregs maintain epithelial balance during alveolar development

The alveolar epithelium is composed of two principal cell types: alveolar type I (AT1) cells, which are elongated and specialised for gas exchange, and alveolar type II (AT2) cells, which produce surfactant and act as progenitors for epithelial regeneration^15^. In LPS-induced adult acute lung injury models, Treg depletion leads to upregulation of interferon (IFN) signalling in AT2 cells from the injured lung^16^. Consistently, transwell co-culture *in vitro* experiments demonstrate that Tregs directly promote AT2 proliferation in a contact-independent manner^12^. Together, these findings suggest Tregs actively regulate epithelial cell function and regenerative capacity. Given that alveologenesis in early life requires tightly coordinated epithelial differentiation and remodelling, we hypothesised that neonatal lung Tregs may play a critical role in regulating alveolar development by controlling inflammatory signals and supporting epithelial maturation.

To test this, we used Foxp3-DTR mice to selectively deplete Tregs during the first week of life. With PBS-treated mice as controls, diphtheria toxin (DT) was administered at P1, P2, and P4 (Fig. 2A), resulting in efficient depletion of neonatal Tregs at P5 (ΔneoTreg; SFig2A). Tregs in both the lung and lymph nodes gradually recovered over time, reaching levels above PBS control by P22 (SFig2B).

**Figure 2.**
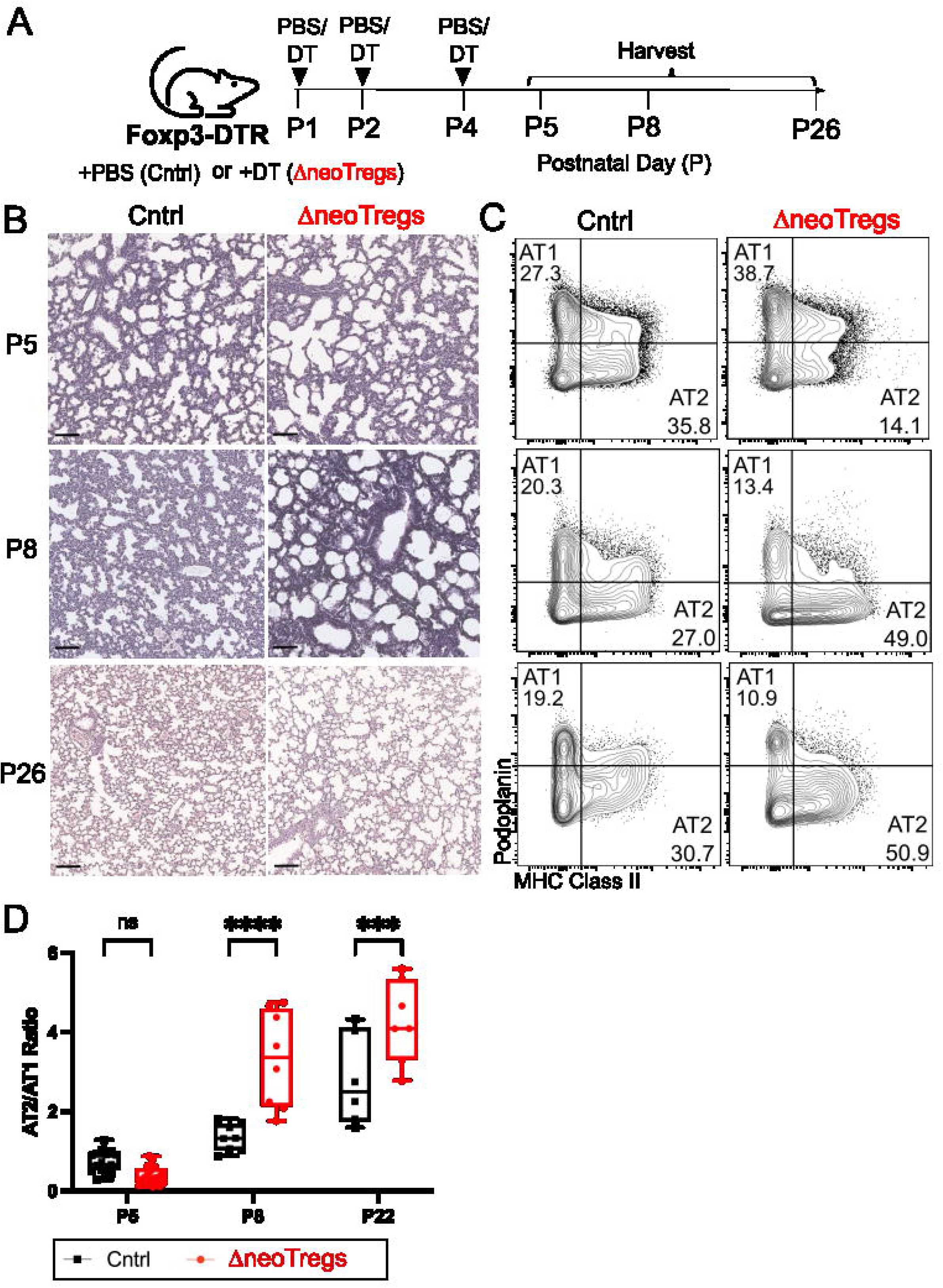
Neonatal Treg depletion disrupts alveolar epithelial maturation during postnatal lung development. (A) Experimental schematic of Foxp3-DTR mice treated with PBS (control) or diphtheria toxin (DT) to induce neonatal Treg depletion (ΔneoTregs). DT or PBS was administered at postnatal days P1, P2, and P4, and lungs were harvested at the indicated time points (P5, P8, and P26). (B) Representative H&E stained lung sections from control and ΔneoTregs mice at P5, P8, and P26. Scale bars as indicated. (C) Representative flow plot and (D) quantification of alveolar epithelial cell populations: AT1 (Podoplanin^+^MHCII^−^) and AT2 (Podoplanin^−^MHCII^+^) populations in control and ΔneoTregs mice at P5, P8, and P26. Data are presented as mean ± SEM. Each point represents an individual sample, pooled from 4 experiments with n =6-10 for each condition. Statistical significance was determined using one-way ANOVA. ns, not significant; ***p < 0.001; ****p < 0.0001.

Histological analysis using haematoxylin and eosin (H&E) staining revealed that Treg depletion led to marked disruption of lung architecture at later time points (Fig2B). While no overt structural differences were observed immediately after depletion at P5, ΔneoTreg lungs displayed enlarged airspaces and impaired septation by the early alveolar stage (P8), indicative of defective alveologenesis. These abnormalities were further exacerbated at the late alveolar stage (P26), demonstrating that transient neonatal Treg loss results in persistent defects in lung structure.

To further characterise epithelial composition in the developing lung, we quantified PDPN^+^MHCII^−^ alveolar type 1 (AT1) and PDPN^−^MHCII^+^ alveolar type 2 (AT2) cells using flow cytometry based on previous studies^12,17^ (Gating strategy in SFig1B). Consistent with histological observations, epithelial balance was comparable between control and Treg-depleted lungs at P5 (Fig2C&D). However, ΔneoTreg lungs exhibited a significant increase in the AT2/AT1 ratio at P8. This imbalance was further amplified at P26, indicating a sustained skewing of epithelial composition toward AT2 cells. Collectively, these findings demonstrate that neonatal Tregs are required to maintain alveolar epithelial cell balance, and subsequentially proper alveolar structure formation.

### Neonatal Tregs restrain inflammatory programmes and neutrophil accumulation

To investigate the mechanisms underlying Treg-driven lung development, we performed whole tissue bulk RNAseq at P5, immediately following neonatal Treg depletion. Differential gene expression analysis revealed that Treg-deficient lungs upregulated neutrophil-attracting chemokines (*Cxcl10*) alongside multiple interferon-related inflammatory genes, including *Ifit1, Irgm1* and members of the *Gbp* family (Fig3A&B).

Immunofluorescence staining and flow cytometry confirmed a marked increase in Ly6G^+^ neutrophils in ΔneoTreg lungs at P5 (Fig. 3B–E), indicating enhanced neutrophil accumulation in the absence of neonatal Tregs. Together, these findings demonstrate that neonatal Tregs play a critical role in restraining early-life neutrophilic inflammation.

**Figure 3.**
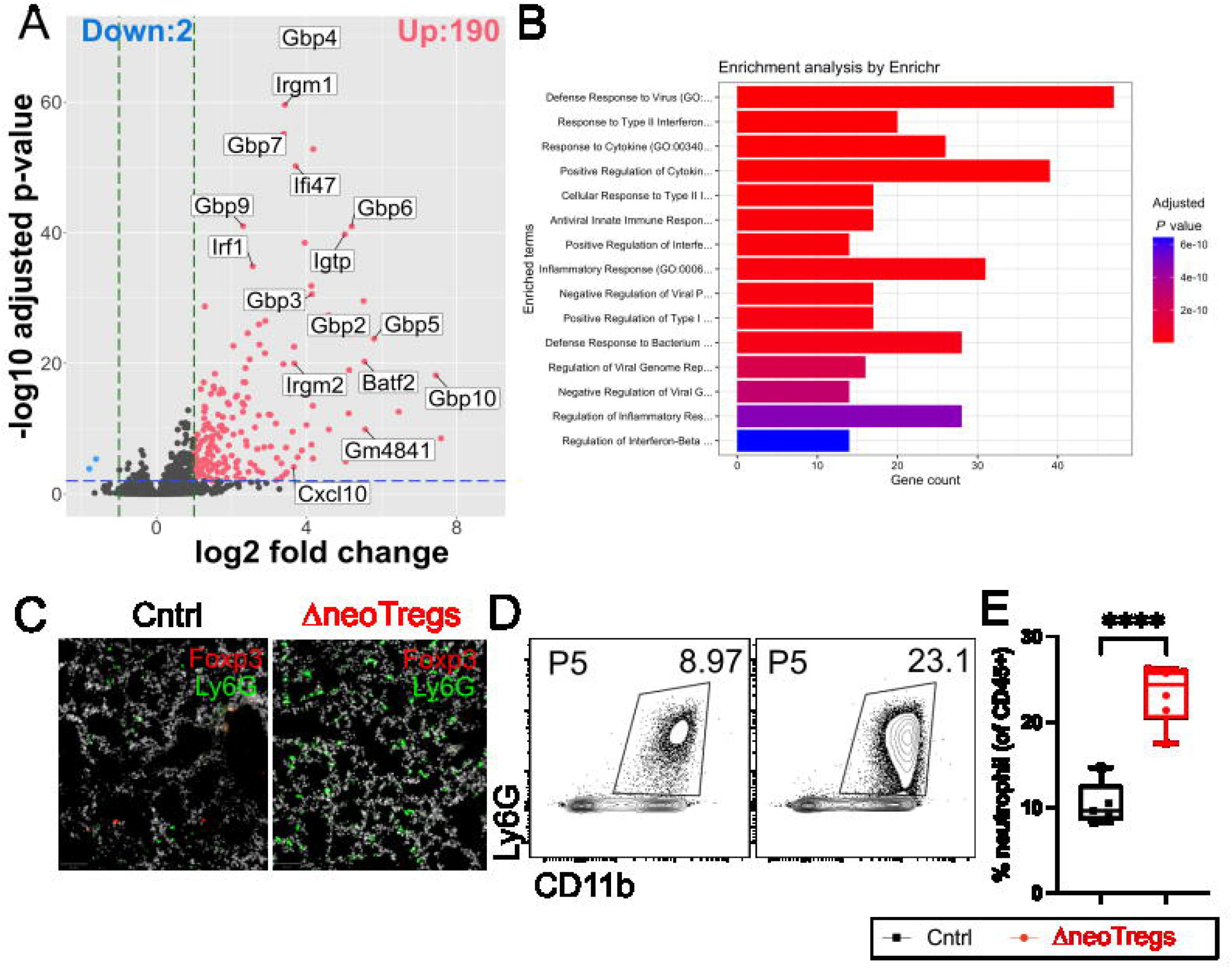
Neonatal Treg depletion induces interferon-driven inflammatory responses and neutrophil accumulation in the lung. (A) Volcano plot of differential gene expression in lungs from control and ΔneoTregs mice. (B) Gene ontology (GO) enrichment analysis of upregulated genes. (C) Representative immunofluorescence images of lung sections from control and ΔneoTregs mice stained for Foxp3 (red) and Ly6G (green). (D) Representative flow cytometry plots and (E)quantification of CD11b+Ly6G+ neutrophils (gated from live CD45+ cells) in Treg sufficient and deficient lung at P5. Data are presented as mean ± SEM. Each point represents an individual sample, pooled from 4 experiments with n = 5-8 for each condition. Statistical significance was determined using unpaired t-test. ****p < 0.0001.

### Neutrophils contribute to the alveolar defects caused by neonatal Treg depletion

To determine whether neutrophils drive the observed long-term lung developmental defects, we combined neonatal Treg ablation with anti-Ly6G–mediated neutrophil depletion and analysed lungs at P26 (Fig4A). Histological quantification of alveolar area showed that neutrophil depletion partially restored alveolar architecture, with reduced airspace enlargement (Fig4B&C). Flow cytometry data further showed the increased AT2 fraction observed following neonatal Treg depletion was attenuated upon neutrophil depletion (Fig4D&E), indicating the restoration of epithelial balance. These data demonstrate that neonatal Tregs promote alveolar development, at least in part, by restricting neutrophil accumulation.

**Figure 4.**
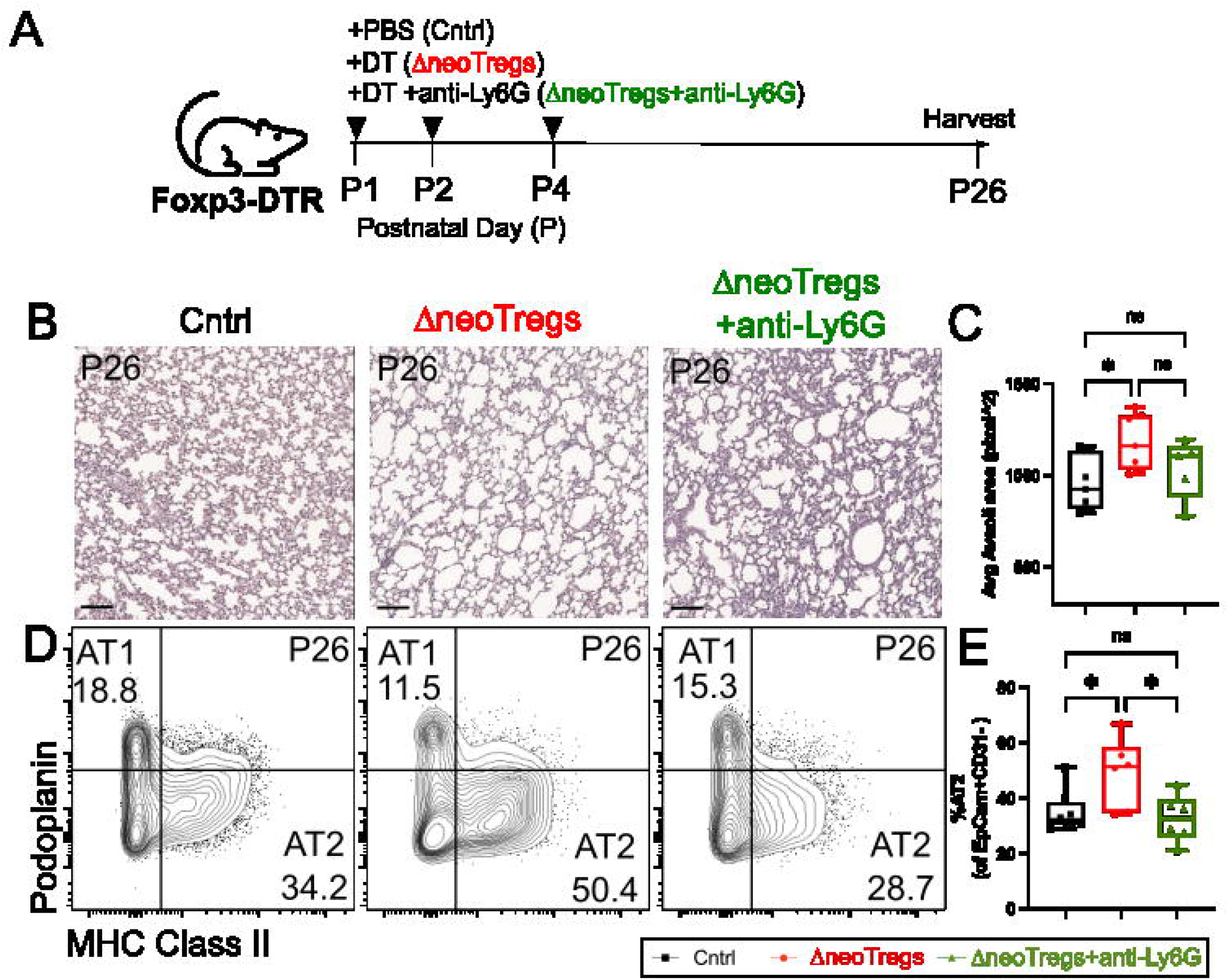
Neutrophil depletion rescues impaired alveologenesis caused by neonatal Treg depletion. (A) Experimental schematic of Foxp3-DTR mice treated with PBS (control), diphtheria toxin (DT; ΔneoTregs), or DT with anti-Ly6G antibody (ΔneoTregs + anti-Ly6G). Treatments were administered at postnatal days P1, P2, and P4, and lungs were harvested at P26. (B) Representative H&E-stained lung sections at P26. Scale bars as indicated. (C) Quantification of average alveolar area from H&E images. (D) Representative flow cytometry plots and (E) quantification of alveolar epithelial cell populations in respond to each treatment. Data are presented as mean ± SEM. Each point represents an individual sample, pooled from 3 experiments with n = 6-7 for each condition. Statistical significance was determined using one-way ANOVA. ns, not significant; *p < 0.05.

Collectively, our findings identify neonatal lung Tregs as key regulators of early-life lung development. During a defined postnatal window around P5, these cells undergo rapid expansion and exhibit a heightened activation state compared to adult Tregs. They function to restrain interferon-driven inflammatory programs and limit neutrophil accumulation within the developing lung. In their absence, inflammatory dysregulation is accompanied by disrupted epithelial balance and impaired alveolarisation, resulting in persistent structural abnormalities. Together, these findings support a role for neonatal Tregs in coordinating immune regulation during postnatal lung development.

## Discussion

The early neonatal period appears to be critical for establishment of the lung Treg compartment, as lineage-tracing studies have shown that nearly 50% of lung Tregs in 8-week-old mice are seeded within the first two weeks of life^18^. However, despite extensive insight into the functions of adult lung Tregs during acute lung injury, their role in lung development remains largely unexplored.

Here, we identify neonatal lung Tregs as key regulators of alveologenesis. We show that a transient, highly proliferative Treg population emerges during early postnatal life and is required for proper alveolar development. Disruption of Tregs during this narrow developmental window leads to persistent defects in lung architecture and epithelial composition. These findings indicate that early-life immune regulation actively shapes developmental outcomes.

Mechanistically, our data show that neonatal Tregs coordinate alveolar development by restraining inflammatory responses and maintainingalveolar epithelial balance. Treg depletion disrupts the AT2/AT1 balance, leading to accumulation of AT2 cells and reduction of AT1 cells. This is accompanied by increased interferon-related inflammatory signalling and neutrophil accumulation, consistent with the established role of Tregs in limiting inflammation during adult lung injury. Importantly, neutrophil depletion partially rescues both the structural defects and the epithelial imbalance, suggesting that neutrophils act as downstream effectors of Treg-mediated alveologenesis. Notably, transient depletion of Tregs during early life leads to long-lasting structural abnormalities, highlighting a critical developmental window before P5. During this period, immune regulation appears essential, and dysregulated inflammation can have lasting consequences for lung structure.

Neutrophils are among the earliest immune cells to populate the neonatal lung and contribute to pathogen clearance^19^. In adults, however, excessive or sustained neutrophil activity can be detrimental to tissue remodelling during injury resolution. Neutrophils release proteases, reactive oxygen species and pro-inflammatory cytokines, which can cause alveolar damage, increase endothelial permeability, and disrupt the alveolar-capillary barrier^20^. Our findings suggest that neonatal Tregs restrain early neutrophilic inflammation, thereby preserving an environment that supports epithelial maturation.

The mechanisms by which neonatal lung Tregs regulate alveologenesis remain to be defined. Insights from acute lung injury provide a useful framework. In adult lungs, Tregs limit inflammation and promote epithelial repair through secretion of pro-regenerative factors such as amphiregulin (Areg)^21,22^ and keratinocyte growth factor (KGF)^23^. These mediators enhance AT2 proliferation and support barrier restoration. Tregs can also directly stimulate AT2 proliferation through soluble factors in a contact-independent manner. In addition, they regulate epithelial transcriptional programs by suppressing IFN-driven signalling in AT2 cells^16^. Together, our findings suggest that neonatal Tregs may employ similar mechanisms to balance inflammation and epithelial differentiation during alveologenesis and thereby ensure proper lung development.

In summary, this study demonstrates that neonatal Tregs modulate lung development by restricting neutrophil infiltration and maintaining alveolar epithelial balance. These results reveal a previously unappreciated role for immune regulation in lung development and suggest that disruption of Treg function during early life may have long-term consequences for pulmonary structure and function

## Acknowledgments

We thank the Advanced Cytometry Platform (Flow Core), Research and Development Department at Guy’s and St Thomas’ NHS Foundation Trust, and the Barts Cancer Institute Flow Cytometry Facility at Queen Mary University of London for assistance with flow cytometry experiments.

## Funding

We acknowledge support by the following grant funding bodies: This work was supported by a Sir Henry Dale Fellowship jointly funded by the Wellcome Trust and the Royal Society awarded to N.A (Grant Number 213401/Z/18/Z). J.Z.X and P.P.L. are supported by Wellcome Trust PhD fellowships (218452/Z/19/Z) and (108874/B/15/Z), respectively.

## Author contributions

Conceptualization, N.A.; methodology, formal analysis, N.A., M.S., P.L., H.A.; investigation, M.S., P.L., H.A., J.Z.X.; manuscript preparation, N.A., M.S., P.L.; supervision, N.A.; funding acquisition, N.A.

## SUPPLEMENTARY FIGURE LEGENDS

**Supplementary Figure 1. Flow cytometry gating strategy for lung Treg and alveolar epithelial cells.** Cells were first stained with Zombie UV live/dead stain and CD45 to identify live cells. (A) T cell panel. T cells (CD45+TCRgd-CD3+) populations were further segregated based on CD8 and CD4 expression. Tregs were gated based on their Foxp3 expression. (B) Flow cytometric gating strategy to identify lung alveolar epithelial cell populations. Cells were identified from live CD45neg gate, followed by identifying EpCAM+ CD31-epithelial cells. Within the epithelial compartment, alveolar type 1 (AT1) cells were defined as PDPN^+^MHCII^−^, and alveolar type 2 (AT2) cells as PDPN^−^MHCII^+^.

**Supplementary Figure 2. Efficient Treg depletion and recovery kinetics following neonatal ablation.** (A) Representative flow cytometry plots and quantification of Tregs in Treg-depleted Foxp3 DTR mice. Kinetics of Treg recovery in the (B) lung and (C) lymph nodes following neonatal depletion at indicated time points. Data are presented as mean ± SEM. Each point represents an individual sample, pooled from 3 experiments with n = 6 for each condition. Statistical significance was determined using unpaired t-test. ****p < 0.0001.

## Methods

### Animal study design

All animal experiments were conducted in compliance with the UK Animals (Scientific Procedures) Act 1986 under a UK Home Office licence (PP6051479) and with approval from the King’s College London ethical review committee (PP70/8474; establishment licence X24D82DFF). Studies were reported in accordance with ARRIVE guidelines. Only procedure-naïve animals were used, and all efforts were taken to minimise distress. Animals were sacrificed through overdosage of anaesthesia, confirmed with circulation cessation.

*Foxp3*^*tm3(HBEGF/GFP)Ayr*^/J (Foxp3-DTR, JAX #016958) were purchased from Jackson Laboratory. Both male and female mice aged postnatal day (P)1 to P43 were used for experiments. For Treg depletion, Foxp3-DTR mice were injected intraperitoneally at 50ng/g of diphtheria toxin (DT) on P1, P2 and P4, and harvested on indicated timepoint. For neutrophil depletion, 20-50 μg/g of rat anti-mouse Anti-Ly6G (clone: 1A8; BE0075-1-25MG, 2BScientific) were administrated intraperitoneally on P1, P2, P4 and P6, before sacrificed on P8 for histological and cellular analysis.

### Tissue Processing

Euthanised mice were subjected to a small incision in the right ventricle and the pulmonary circulation was perfused by steadily injecting 20ml ice-cold PBS into the left ventricle. Lungs were carefully dissected and lobes were separated.

For histology, lungs were fixed in 4% Paraformaldehyde (PFA) for 4h at 4°C, washed with PBS before transferring to 30% sucrose for OCT embedding or 70% ethanol for paraffin embedding.

For flow cytometry, lung lobes were finely chopped and incubated in 5 ml enzyme R10 solution (10% Fetal Bovine Serum, 1% penicillin-streptomycin (P4458, Sigma-Aldrich) in RPMI-1640 with L-glutamine) supplemented with collagenase A (1 mg/mL; 1010357800, Roche) and DNAse1 (0.1 mg/mL; 10104159001, Roche). The lobes were digested for 1 hour at 37 °C with rotation and the reaction was quenched with double the amount of ice cold R10 media. The samples were filtered through a 70 μm sterile filter and the cell pellet was resuspended in Ammonium-Chloride-Potassium (ACK) red blood cells lysis buffer (A10492-01, Gibco) for 1 min at room temperature. The reaction was quenched with 20 ml R10 media, pelleted, resuspended in 500 μl of FACS buffer (2% FBS, 2 mM EDTA in PBS) and cells were counted using Nucleocounter (Chemometec).

Single cell suspension were aliquoted in a round bottom 96 well plate and stained with live dead stain (Zombie UVTM, 423107, Biolegend) for 20 min on ice, washed with FACS buffer before stained with surface antibody markers (see Table 1). Cells were washed with FACS buffer and the pellet was fixed permeabilised for 20 min on ice using Foxp3/ Transcription factor Staining buffer kit (eBioscience). The cells were then washed with 1X permeabilization buffer (eBioscience) and the pellet was resuspended with the intracellular antibodies (see Table 1) and incubated on ice for 20 min. Stained cells were run on Fortessa LSRII, Fortessa LSRI and FACSSymphony (BD Bioscience) in KCL BRC Flow Cytometry Core. SPHERO Rainbow calibration particle, 8 peaks beads (559123, BD Biosciences) were used to standardise experiments. UltraComp eBeads™ (01-2222-42, Thermofisher), ArC™ Amine Reactive Compensation Bead Kit (A10346, Thermofisher) were used for compensation. All analysis was performed using FlowJo v10.

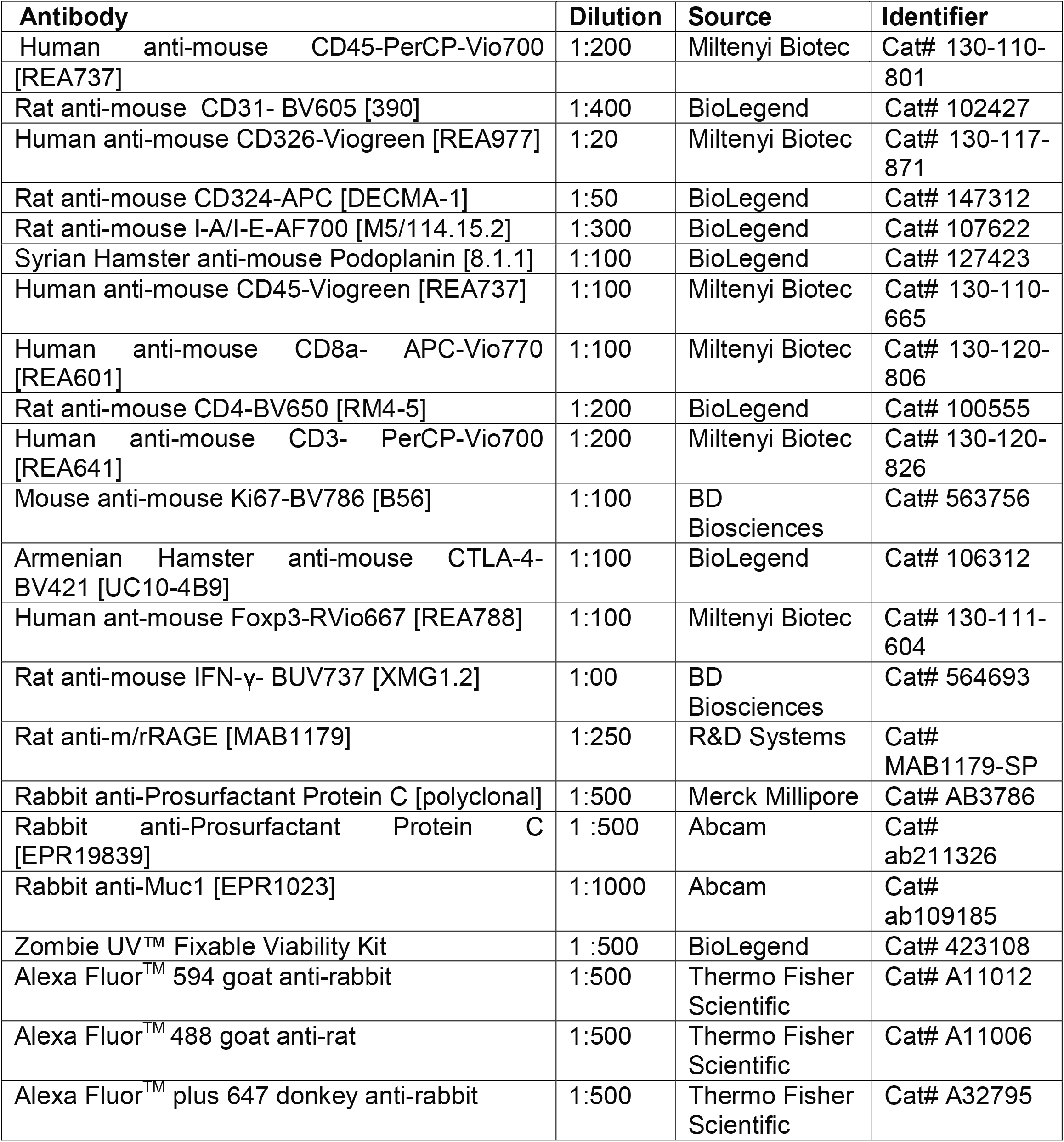

### Immunostaining

For OCT embedding, lungs were fixed in 4% Paraformaldehyde (PFA) for 4h at 4°C, followed by washing in PBS before being embedded in OCT mounting media (361603E, VWR). Staining was performed on 10 μm OCT-embedded sections. Briefly, sections were incubated for 10 min in 0.3% Triton-X-100 (T9284-1L, Sigma-Aldrich) in PBS solution with gentle agitation, before blocking with 1% BSA (A7906-100G, Sigma-Aldrich), 5% Normal Goat Serum (NGS) (31873, Thermo Fisher Scientific) and 0.1% Triton-X-100 for 60 min at room temperature. Sections were then incubated with primary antibodies (see Table 1) overnight at 4°C. The next day, sections were incubated with Alexa Fluor-coupled secondary antibodies (see Table 1) for 2 h at room temperature. DAPI was added to the slides for nuclei staining, following by mounting slides with ProLongTM Gold antifade reagent (P36930, Thermo Fisher Scientific). For H&E staining, lung lobes were initially fixed in 10% neutral-buffered formalin overnight at 4°C, followed by transfer to 70% Ethanol at 4°C. Samples were embedded in paraffin, sectioned at 6 μm and H&E stained according to manufacturer’s instructions (Abcam).

### Image acquisition and analysis

Images were captured at x40 objective using Leica SP8 confocal microscope and lineage-traced cells were quantified manually from 2-3 images per sample using ImageJ. Muc1 domains were quantified manually on pro-SFTPC+ cells enfacing the alveolar lumen using Imaris software. For H&E analysis, images were acquired using NanoZoomer 2.0RS Digital Slide Scanner (Hamamatsu Photonics K.K.) and analysed using NPD viewer software (Hamamatsu, v2.7.43). Protocol by Pua et al., (2005) was adapted to measure alveolar wall thickness was measured using NPD viewer software. Briefly, within 1 μm diameter circle positioned on the image, 3 identically sized squares were placed on the image to represent the regions of interest (ROIs). The thinnest portion from each alveolar wall was measured by measuring the thickness of the wall. Protocol by Koc et al., (2022) was adapted for RAC measurement, images were analysed using ZEN 3.5 (blue edition) where a perpendicular line was drawn from one terminal respiratory bronchiole to the nearest pleural surface. The number of alveoli that crossed the perpendicular line was manually quantified.

### Quantitative PCR (qPCR)

cDNA from lung RNA was synthesised using iScript cDNA synthesis kit (1708891, Bio-Rad) and qPCR was carried out using TaqMan™ PreAmp Master Mix Kit (4384267, ThermoFisher) according to manufacturer’s instructions. Gapdh (Mm99999915_g1, VIC), predicted gene 12250 (Mm01352113_m1, FAM), guanylate-binding protein 10 (Mm03647514_uH, FAM), predicted gene 4951 (Mm01621208_s1, FAM), guanylate binding protein 5 (Mm00463735_m1, FAM), basic leucine zipper transcription factor, ATF-like 2 (Mm01231591_m1, FAM) and interferon gamma induced GTPase (Mm01347794_g1, FAM) TaqMan probes were used to identify genes of interest. Samples were run in triplicate on CFX384 Touch Real-Time PCR system (BioRad).

### Whole tissue total RNA isolation, bulk-RNA sequencing and analysis

Post-caval lung lobes were preserved in RNAlater (Sigma; R0901-500ML) at -80°C. Tissues were defrosted and homogenised using Precellys® tubes containing 1.4 mm ceramic beads (432-3751, VWR) in RLT lysis buffer (74104, Qiagen). RNA extraction using RNeasy Mini Kit (50) (74104, Qiagen) and DNase incubation using RNase-Free DNase Set (50) (79254, Qiagen) was performed according to manufacturer’s instructions. RNA was to sent to Genewiz (AZENTA Life Sciences) for Illumina NovaSeq 2x150 bp paired-end mRNA sequencing and RNA quality number (RQN) was assessed using Fragment Analyzer. Differential gene expression analysis was performed on R software using DEseq2, ggplot2 and EnhancedVolcano package.

### Statistical analysis

Statistical comparisons for two independent groups was performed using two-tailed unpaired t-test or Mann Whitney t-test. Comparisons between three or more independent groups was analysed using Ordinary One-way ANOVA with Tukey’s multiple comparisons test. Comparisons of more than one variable between three or more independent groups was performed using Two-way ANOVA with Tukey’s multiple comparisons tests. *p < 0.05, **p < 0.01 ***p < 0.001, ****p < 0.001, ns >0.05.

## Notes

### Competing Interest Statement

The authors have declared no competing interest.

