## Supplementary figures and images for "Neonatal regulatory T cells promote alveologenesis by restraining neutrophil-mediated inflammation"

### SUPPLEMENTARY FIGURE 1

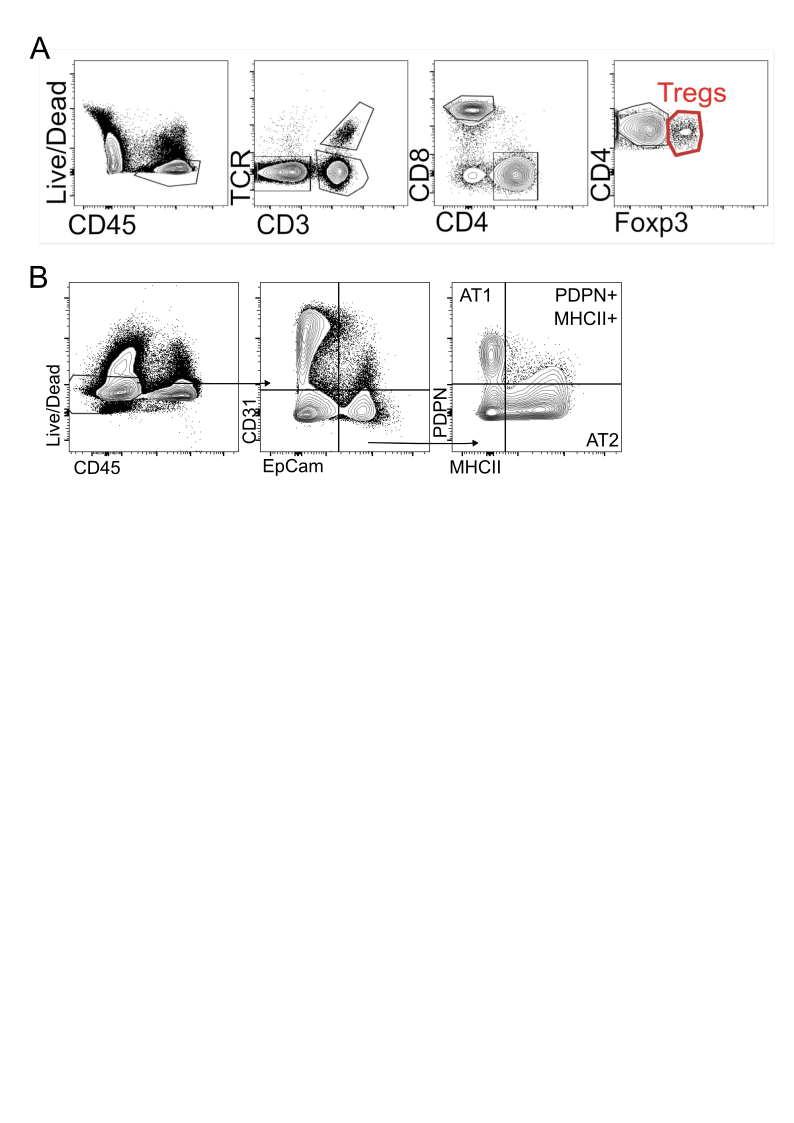

### SUPPLEMENTARY FIGURE 2

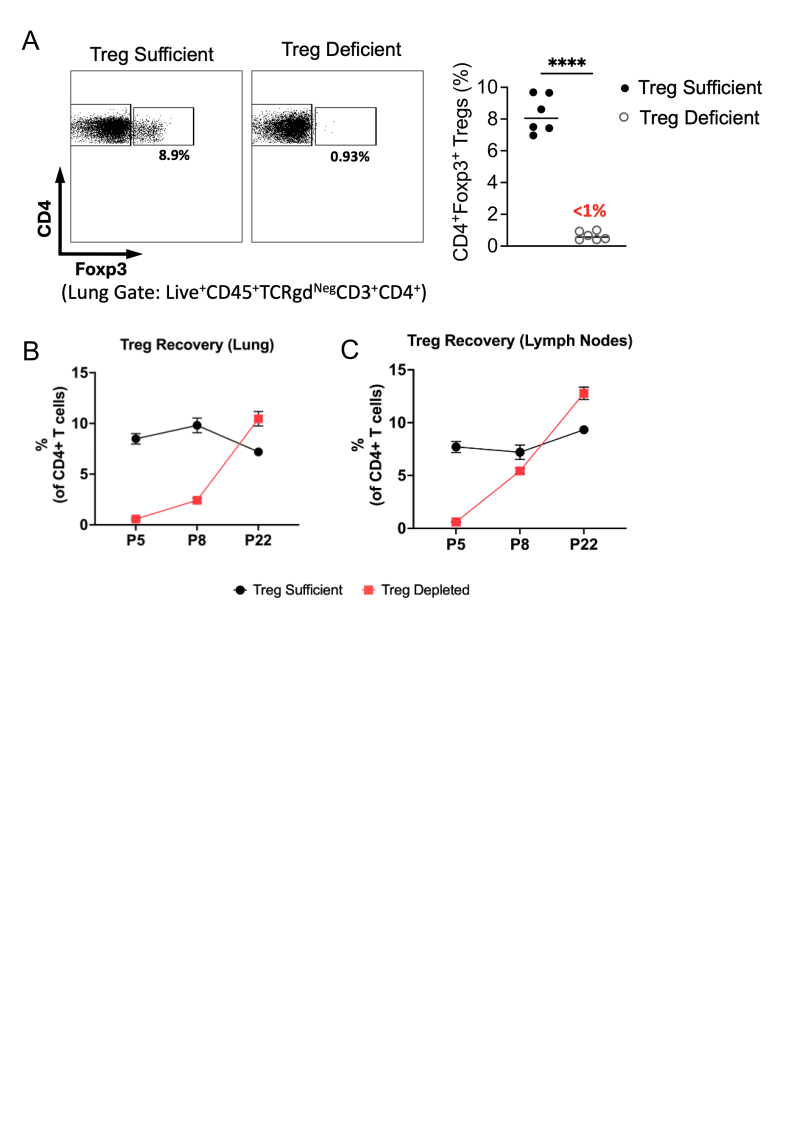
